# Erosion of human X chromosome inactivation causes major remodelling of the iPSC proteome

**DOI:** 10.1101/2020.03.18.997049

**Authors:** Alejandro J. Brenes, Harunori Yoshikawa, Dalila Bensaddek, Bogdan Mirauta, Daniel Seaton, Jens L. Hukelmann, Hao Jiang, Oliver Stegle, Angus I. Lamond

## Abstract

X chromosome inactivation (XCI) is a dosage compensation mechanism in female mammals whereby genes from one X chromosome are repressed. Analysis of human induced pluripotent stem cell (iPSC) lines using proteomics, RNAseq and polysome profiling showed a major change in the proteome upon XCI erosion. This resulted in amplified RNA and protein expression from X-linked genes. However, increased protein expression was also detected from autosomal genes without a corresponding mRNA increase, altering the protein-RNA correlation between genes on the X chromosome and autosomes. Eroded iPSC lines display ~13% increase in cell protein content, along with increased expression of ribosomal proteins, ribosome biogenesis and translation factors. They also showed significantly increased levels of active polysomes within the eroded lines. We conclude that erosion of XCI causes a major remodelling of the proteome, with translational mechanisms affecting the expression of a much wider range of proteins and disease-linked loci than previously realised.

## Introduction

In humans and other mammalian species, female cells have two copies of the X chromosome, while males have a single X chromosome and a much smaller Y chromosome that is not present in females. The structure of one of the two female X chromosomes is altered, causing repression of transcription and thereby inactivating expression of alleles located on this second copy of the X. This process is termed X chromosome inactivation (XCI).

The XCI process in female cells is considered to be a critical dosage compensation mechanism that evolved in mammals as a way to equalize X-linked gene expression (1, 2). XCI is vital for embryonic development and failure to induce XCI has been shown to cause embryonic lethality (3, 4). The initiation of XCI is controlled by a specific locus, termed the X-inactivation center (5) (Xic). The mechanism of XCI involves a profound structural reorganization of the inactivated copy of the X chromosome, which becomes transcriptionally silent and heterochromatic (5, 6).

Within the Xic a long non-coding RNA, called ‘XIST’, has been shown to be a vital component of the XCI process (7, 8). Accumulation of XIST RNA across the inactive copy of the X chromosome triggers gene silencing and chromatin changes that produce the transcriptionally inactive state (9, 10). A major clinical consequence of human skewed XCI is the occurrence of gender-specific genetic disorders that result from XCI preventing expression of the single wild type allele for X chromosome linked disease loci in heterozygous females.

In this study, we explore the global consequences for human gene expression when XCI is eroded in female cells. Specifically, we have analysed the impact of XCI erosion using multiple human induced pluripotent stem cell (iPSC) lines that were all reprogrammed from primary skin fibroblasts taken from healthy female donors (11). As many as 40% of these iPSC lines showed low levels of XIST RNA expression and we found this correlated strongly with eroded XCI. We performed a global analysis comparing in parallel RNA and protein expression levels in 74 independent iPSC lines derived from healthy female donors that were stratified based upon having either high, or low levels of XIST RNA expression.

This study provides the first in depth analysis by mass spectrometry-based proteomics of how erosion of XCI affects gene expression in human cells at the protein level. All of the raw and processed MS files have been deposited to PRIDE (12) [PXD010557], while the processed, protein level data have been integrated into the Encyclopedia of Proteome Dynamics (13), providing a convenient open access online database for interactive data exploration.

The data show that erosion of XCI in human iPSCs activates both transcription and increased protein production from genes on the inactive X chromosome. Remarkably, we also uncover a major increase in the abundance of many proteins encoded by genes on the autosomes, independent of a parallel increase in transcription. Female cells with low XIST RNA show a median ~13% increase in total protein levels, along with higher levels of polysomes and components of the translational machinery. These data indicate that erosion of XCI can affect the expression of a much wider range of proteins and disease-linked gene loci than previously realised based on RNA analysis alone.

## Results

### RNAseq and proteomic datasets

All of the iPSC lines used for this study were reprogrammed from human primary skin fibroblasts as previously described (11). The mass spectrometry-based proteomic analysis was performed using tandem mass tagging (TMT) (see Methods; Fig.1a). This study analyses data from 74 independent iPSC lines derived from healthy female donors with matching proteomics and RNAseq data generated from the same cell lines. In total, we detected expression of >9,500 protein groups (protein isoforms without discriminating peptides; hereon proteins) at 1% FDR, with a median of 8,479 proteins identified across all lines (Fig. 1b) and with a median protein sequence coverage of 42% across all proteins (Fig. 1c). All downstream analysis was performed on a subset of proteins (8,908) identified with at least 3 ‘Razor + unique peptides’ (see methods). To compare protein expression levels between the respective lines, protein copy numbers were estimated using the ‘proteomic ruler’ approach (14, 15). This is well suited for the analysis of HipSci lines, which have been shown to have near identical DNA content.

**Figure 1.**
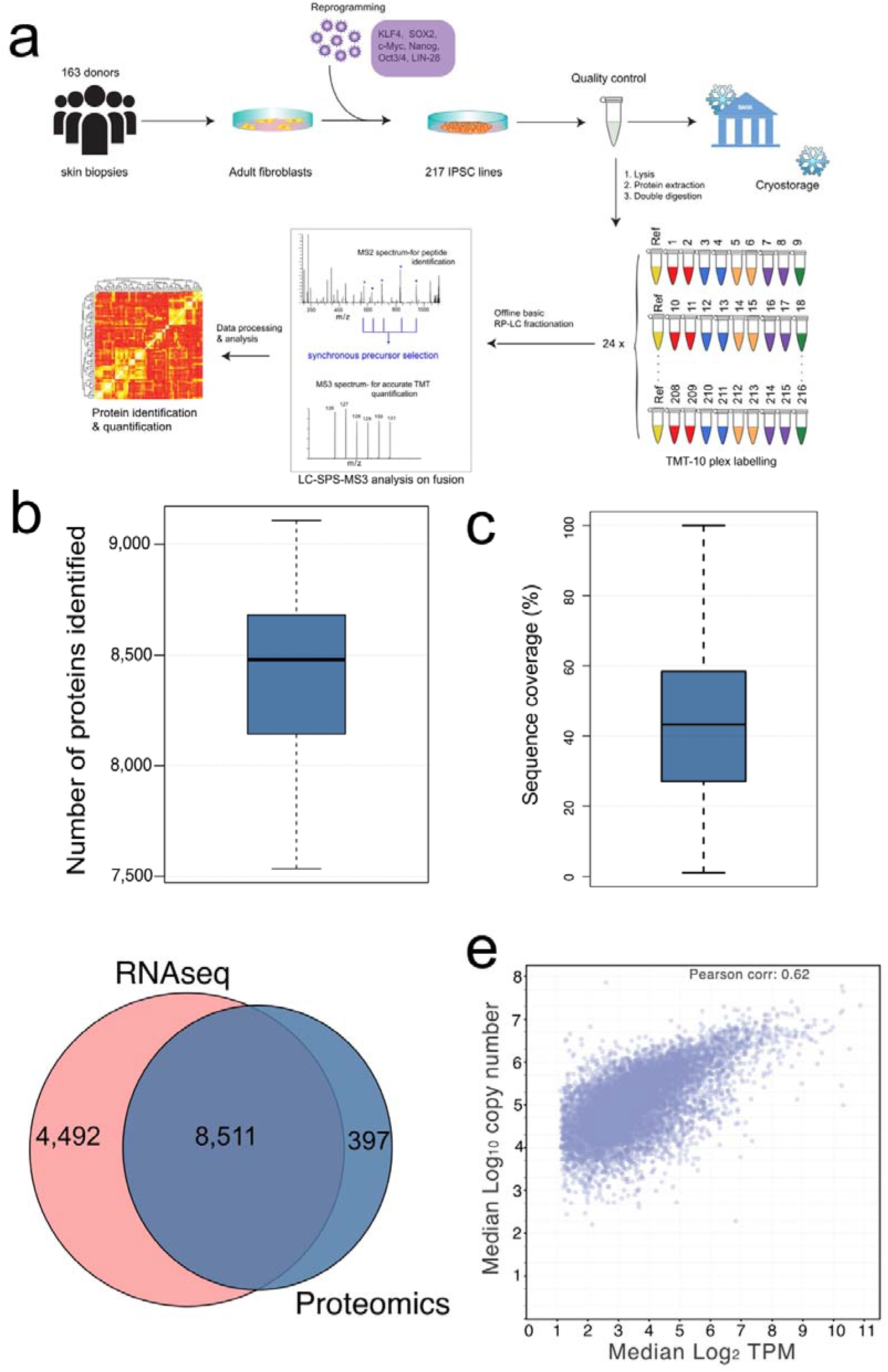
Comprehensive coverage: **(a)** The HipSci proteomics workflow from reprogramming to identification and quantification. **(b)** Boxplot showing the number of proteins identified per line across the 66 filtered (see methods) female iPSC lines. The lower and upper hinges represent the 1st and 3rd quartiles. The upper whisker extends from the hinge to the largest value no further than 1.5 * IQR from the hinge, the lower whisker extends from the hinge to the smallest value at most 1.5 * IQR of the hinge. **(c)** Boxplot showing the sequence coverage for all proteins detected within the dataset. The lower and upper hinges represent the 1st and 3rd quartiles. The upper whisker extends from the hinge to the largest value no further than 1.5 * IQR from the hinge, the lower whisker extends from the hinge to the smallest value at most 1.5 * IQR of the hinge. **(d)** Pie chart showing the overlap between quantified gene products in the proteomics and RNAseq datasets. **(e)** Scatter plot showing the median log_2_ transcripts per million vs the median log_10_ copy number for all gene products.

From the RNAseq data, a total of 12,798 transcripts were quantified after filtering (see methods), from which there was evidence from mass spectrometry of protein expression for 65% of these transcripts (Fig. 1d). To explore the relationship between RNA and protein abundance levels in this set of iPSC lines, the Pearson correlation of the median log_2_ transcripts per million (TPM) vs the log_10_ protein copy numbers was calculated (Fig. 1e). The resulting Pearson correlation coefficient of 0.62 is not unexpected as it is similar to what has been determined in multiple previous studies for different human cell types and for other mammalian species (16–18).

### X-chromosome inactivation

There is previous evidence that some human primed iPSCs can exhibit erosion of XCI, where the X chromosome loses H3K27me3 marks as well as XIST RNA expression (19, 20). Although the role of XIST, a long non-coding RNA molecule that covers the inactive copy of the X chromosome (21), remains unclear, along with its relation to escape from XCI, it has been reported that loss of XIST expression is characteristic of class III hESCs that display eroded XCI (22). We evaluated the correlation between levels of XIST RNA expression and erosion of XCI within the HipSci iPSC lines. As a measure of eroded XCI, we used the RNAseq data to detect biallelic expression from X-linked genes (Fig. 2a). This showed a clear correlation between iPSC lines showing low XIST RNA expression and increased levels of biallelic expression for X-linked genes. Interestingly, a parallel comparison of biallelic expression and levels of expression of XACT RNA, another long non-coding RNA implicated in the mechanism of XCI in human(23), showed little or no correlation with increased levels of biallelic expression from the X chromosome in these iPSC lines (supplemental figure 1). We conclude, therefore, that reduced levels of XIST RNA expression provides a good marker for erosion of XCI in this cohort of iPSC lines, while expression of XACT RNA does not.

**Figure 2.**
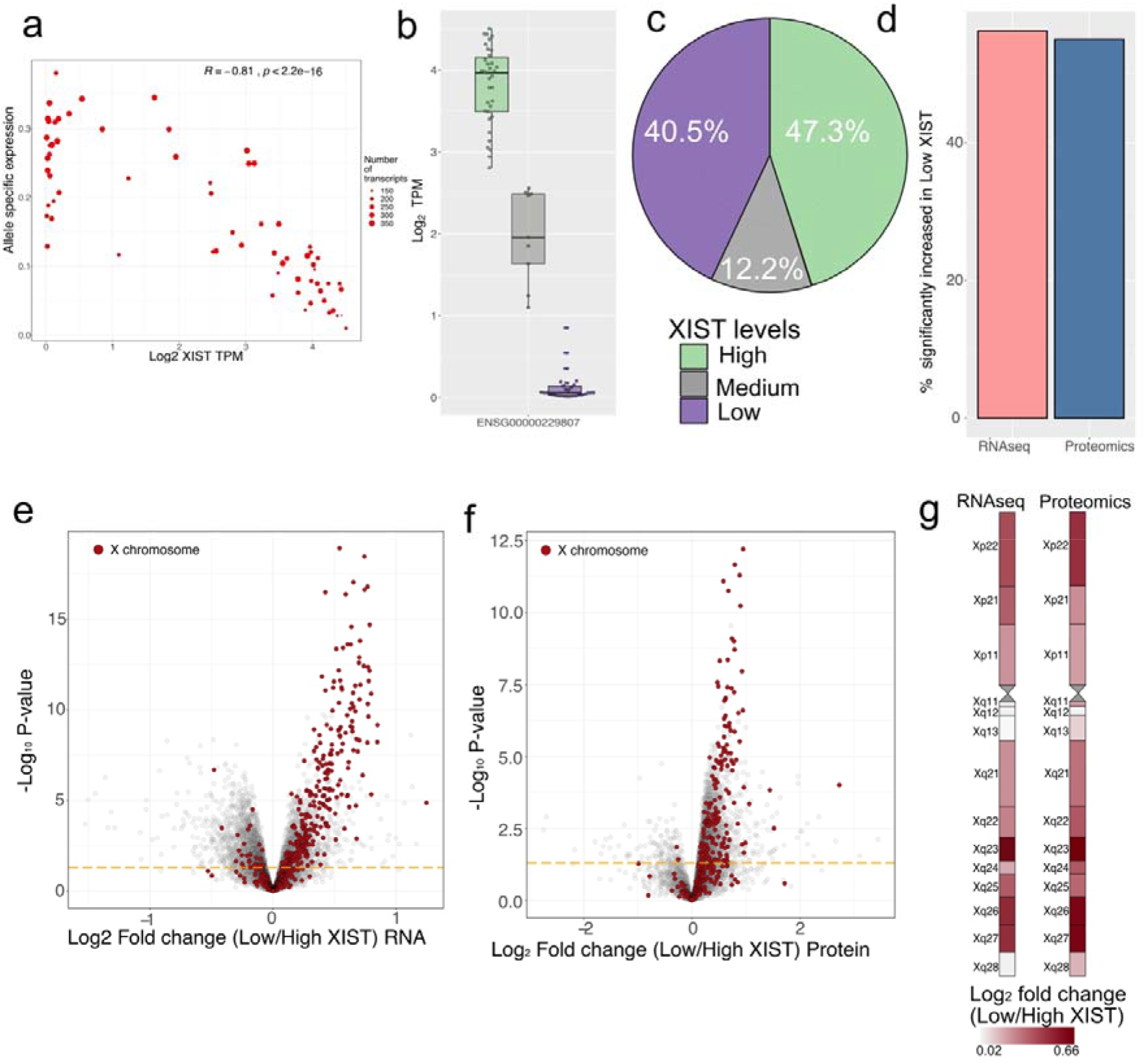
XIST and XCI: **(a)** Scatter plot showing the allele specific expression (ASE) for all X-linked transcripts vs the log_2_ XIST TPM for all 74 healthy female lines. The size is determined by the number of transcripts used for the ASE analysis **(b)** Box plot showing Log_2_ Transcripts per million (TPM) for the long non-coding RNA XIST across all 3 populations; Low, Medium and High. The lower and upper hinges represent the 1st and 3rd quartiles. The upper whisker extends from the hinge to the largest value no further than 1.5 * IQR from the hinge, the lower whisker extends from the hinge to the smallest value at most 1.5 * IQR of the hinge. **(c)** Pie chart showing the percentage of healthy female lines within each XIST stratified population. **(d)** Bar plot showing the percentage of X chromosome gene products which are significantly increased in expression (q-value<0.05) for both RNAseq and proteomics datasets **(e)** Volcano plot showing the RNA log_2_ fold change between the Low and High XIST populations on the X axis, with the −log_10_ p-value on the y axis. X chromosome transcripts are highlighted in red; autosome transcripts are coloured grey. All transcripts above the orange line have a p-value lower than 0.05 (**f)** Volcano plot showing the protein log_2_ fold change between the Low and High XIST populations on the X axis, with the −log_10_ p-value on the y axis. X chromosome proteins are highlighted in red; autosomal proteins are coloured grey. All proteins above the orange line have a p-value lower than 0.05 **(g)** X chromosome map showing the median log_2_ fold change (Low/High XIST) across chromosomal bands for both the RNAseq and proteomic datasets.

We stratified the 74 female iPSC lines into Low, Medium and High XIST RNA populations, based on the RNAseq expression data (Fig. 2b). This showed that two main populations were distinguished, with 40.5% of the iPSC lines having very low levels of XIST RNA expression, in comparison with 47.3% showing the expected, high levels of XIST RNA (Fig. 2c). We also identified a minor population (12.2%) that showed an intermediate level of XIST expression. Further analysis was focussed on comparing gene expression between the High and Low populations stratified by levels of XIST RNA expression. To improve accuracy, a subset of the High XIST RNA lines were filtered out from subsequent analysis due to technical reasons (see methods), leaving a final comparison focussed on 26 independent iPSC lines expressing High levels of XIST RNA vs 30 independent lines expressing Low levels of XIST RNA.

Next, we analysed RNA and protein expression specifically for genes on the X chromosome, comparing their expression between the respective Low and High XIST RNA populations. This showed ~55% of these X-linked genes having significantly increased expression in the Low XIST population, at both the RNA and protein levels (supplemental table 1&2; Fig. 2d-f).

We mapped the 283 X-linked genes whose expression was quantified in both the RNAseq and proteomic data back to their chromosomal bands and calculated the median log_2_ fold changes across all bands (Fig. 2g). This showed that both the transcriptomic and proteomic data displayed very similar fold change patterns across the X chromosome. This RNA-protein concordance is consistent with a predominantly transcription driven regulation of X-linked genes and further indicates that erosion of XCI is not uniform along the X chromosome. For example, the data show specific locations with higher erosion, such as Xq23. This is consistent with previous observations showing that there are specific loci that can preferentially escape XCI(24, 25). In summary, these data show that erosion of XCI in human cells results in both increased transcription and protein expression for X-linked genes.

### Impact of XCI erosion on Autosomes

We next examined whether the erosion of XCI also affects expression of genes located on the autosomes. For this, we compared the RNAseq and proteomic data for differential expression between the XIST RNA-stratified populations for genes mapped to each individual chromosome. Specifically, we aimed to determine to what degree expression differences detected at the RNA level were also seen at the protein level. We note that the correlation between RNA and protein levels can be compared in two different ways(26, 27). First, by correlating the abundance of transcripts with their corresponding protein products, which here generated a Pearson correlation of 0.62, as described above (Figure 1e). Second, by correlating how closely changes in expression of transcripts and changes in expression of proteins each vary between different conditions. For this latter analysis we calculated the Pearson correlation coefficient for the Log_2_ fold changes in RNA and protein levels between the respective Low vs High XIST RNA populations (Fig. 3a). For all genes detected, the global Pearson correlation coefficient comparing the RNA and protein fold changes, was 0.31. This correlation coefficient is thus much lower for the correspondence between changing levels of transcripts and proteins than the correlation in total RNA and protein abundance. We conclude that the changes in protein expression between the two XIST RNA-stratified populations are not well predicted by RNAseq data alone.

**Figure 3.**
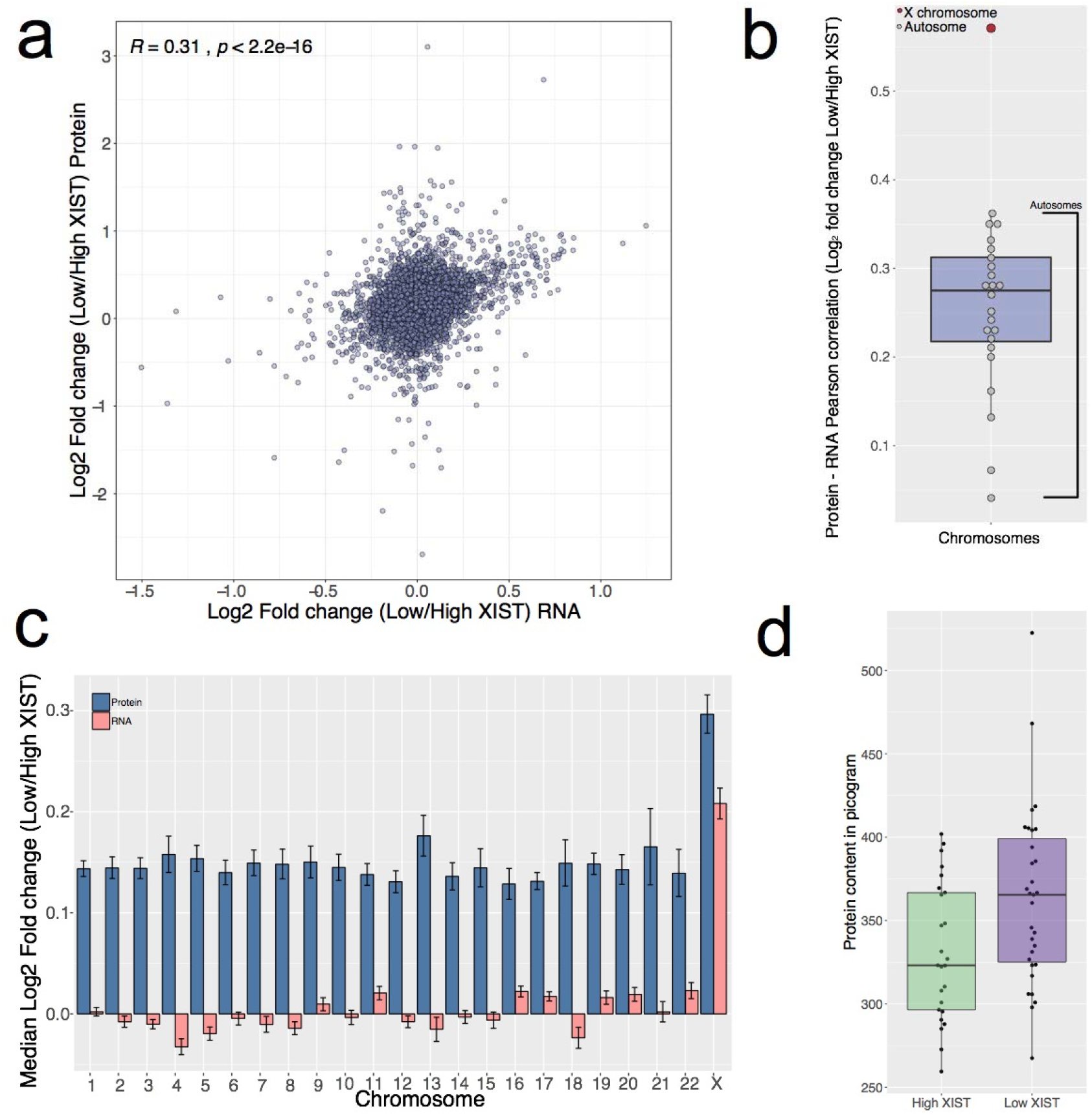
Multi-omic overview: **(a)** Scatter plot showing the RNA vs protein Log_2_ fold change (Low/High XIST) for all gene products quantified at both the protein and RNA level. **(b)** Box plot showing the RNA-Protein log_2_ fold change Pearson correlation for all gene products aggregated at the chromosome level. Autosomes are coloured in grey, the X chromosome is coloured in red. The lower and upper hinges represent the 1st and 3rd quartiles. The upper whisker extends from the hinge to the largest value no further than 1.5 * IQR from the hinge, the lower whisker extends from the hinge to the smallest value at most 1.5 * IQR of the hinge. **(c)** Bar plot showing the median Log_2_ fold change (Low/High XIST) for all gene products aggregated at the chromosome level for both RNAseq and proteomics. The error bars represent the SEM. **(d)** Box plot showing the protein content (see methods) for the High and Low XIST populations. The lower and upper hinges represent the 1st and 3rd quartiles. The upper whisker extends from the hinge to the largest value no further than 1.5 * IQR from the hinge, the lower whisker extends from the hinge to the smallest value at most 1.5 * IQR of the hinge.

To explore this further, we next calculated the sets of RNA-protein correlations individually for each chromosome. We note that our analysis above showed there was a consistent response for both RNA and protein expression changes between the High and Low XIST populations for X chromosome-linked genes (Fig. 2g) and similarly the chromosome specific view showed the X-linked gene products with a Pearson correlation of 0.56 (Fig. 3b). However, the median Pearson correlation for all autosomes was much lower, with a median Pearson correlation coefficient of 0.27 across those chromosomes. Thus, the correlation in the responses of both mRNA and protein levels seen for genes on the X chromosome is an outlier (Fig. 3b). These data indicate a mechanistic difference between the response to Low XIST RNA levels for genes on the X chromosome, in comparison with genes on all of the autosomes.

To explore further this difference detected between the X chromosome and autosomes in how RNA and protein expression levels change in response to erosion of XCI, we compared the median fold change for RNA and protein expression from each gene, across all chromosomes, between the two XIST RNA-stratified populations (Fig. 3c). The highest fold change in expression, which is seen at both the RNA and protein levels, occurs for genes on the X chromosome. However, the proteomics data also revealed increased median fold changes across all of the autosomes. Remarkably, in contrast with genes on the X chromosome, increased protein expression from autosomal genes was predominantly not accompanied by changes in RNA expression (Fig. 3c).

These data indicate that erosion of XCI can result in increased expression of proteins from a much wider set of genes than those located only on the X chromosome. Overall, ~ 27% of the proteins quantified (2,429 out of 8,908), showed significantly increased expression in the Low XIST population (Fig. 2f). In contrast, only 1.2% of proteins (108 out of 8,908), were significantly decreased in expression. Therefore, we hypothesised that the Low XIST population may have an overall higher total protein content. To test this idea, we estimated the total protein content for both stratified populations, based upon the MS data (see methods). This showed that iPSC lines with Low XIST RNA expression had a median ~13% higher total protein content, compared to the lines expressing High levels of XIST RNA (Fig. 3d).

### Proteome Response to XCI

We next investigated in more detail how the proteome was altered in the low XIST population and examined which types of proteins and which protein functions showed changes. First, we performed an overrepresentation test, using Panther(28), focussed on proteins that significantly increased expression in the Low XIST RNA population (q-value<0.05), but without a corresponding significant increase in the RNAseq data. The top-level Gene Ontology (GO) terms these proteins were enriched in were ‘ribonucleoprotein complex biogenesis’ and ‘mRNA metabolic process’. We note that the processes relating to these enriched GO terms have the potential to contribute to post-transcriptional mechanisms that could increase total protein expression from a constant amount of mRNA, including genes related to ribosome biogenesis, ribosomes and protein translation. We calculated Pearson correlation coefficient comparing changes in expression at the respective RNA and protein levels between the XIST RNA-stratified populations for the ribosomal and ribosome biogenesis proteins and found them to be particularly low (Fig. 4a). This is consistent with an important role for post-transcriptional mechanisms in regulating protein expression from these genes.

**Figure 4.**
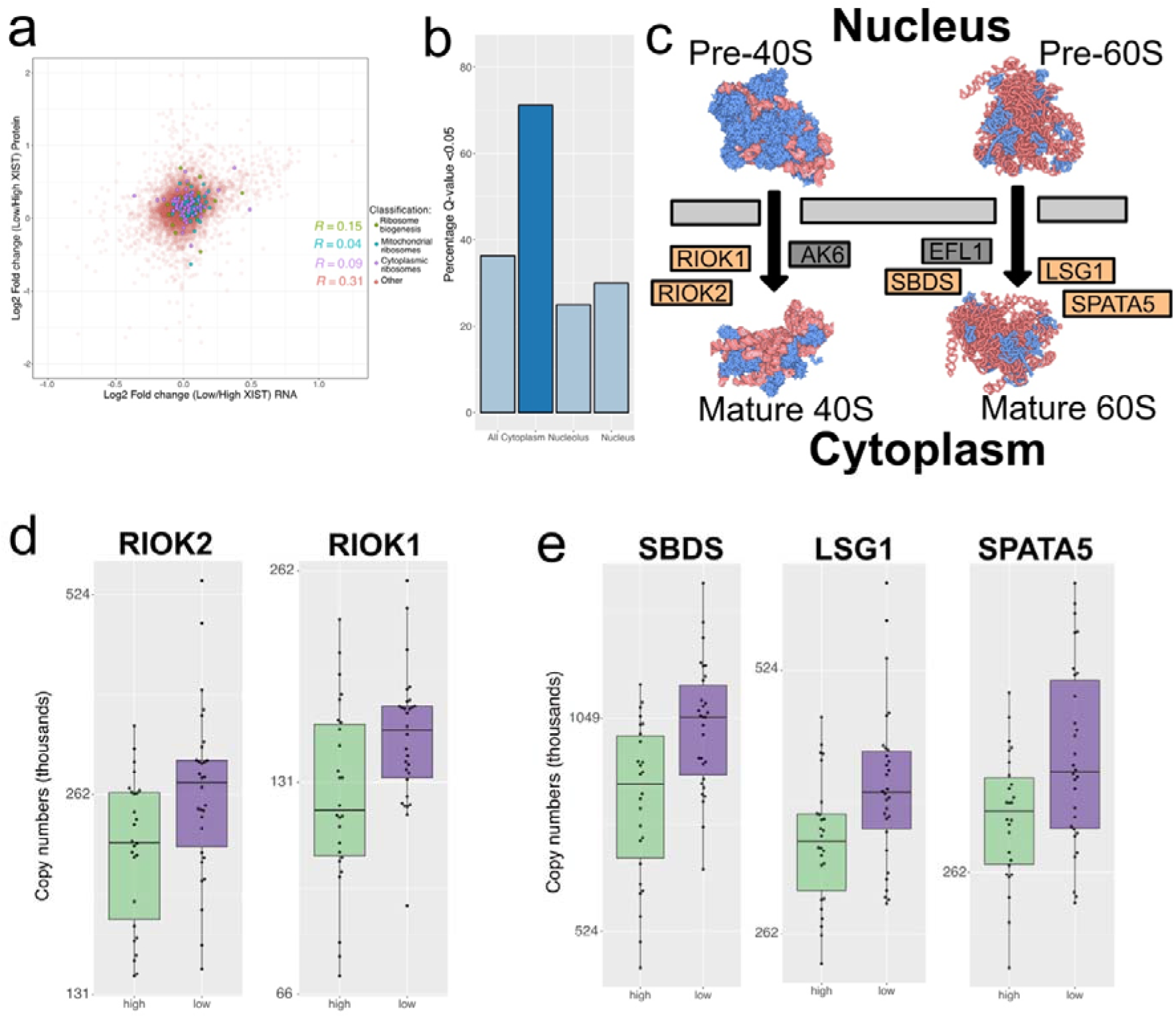
Ribosome biogenesis: **(a)** Scatter plot showing the Log_2_ fold change (Low/High XIST) at the protein and RNA level. Ribosome biogenesis, cytoplasmic and mitochondrial ribosomal proteins are highlighted, and Pearson correlation coefficients provided. **(b)** Bar plot showing the percentage of proteins annotated by Kegg as being part of the ribosome biogenesis pathway which are significantly increased in expression across subcellular localisations **(c)** Schematic showing the cytoplasmic ribosome biogenesis proteins that are significantly increased in expression **(d)** Box plot showing the protein copy numbers for RIOK1 and RIOK2 within the Low and High XIST populations. The lower and upper hinges represent the 1st and 3rd quartiles. The upper whisker extends from the hinge to the largest value no further than 1.5 * IQR from the hinge, the lower whisker extends from the hinge to the smallest value at most 1.5 * IQR of the hinge. **(e)** Box plot showing the protein copy numbers for SBDS, LSG1 and SPATA5 within the Low and High XIST populations. The lower and upper hinges represent the 1st and 3rd quartiles. The upper whisker extends from the hinge to the largest value no further than 1.5 * IQR from the hinge, the lower whisker extends from the hinge to the smallest value at most 1.5 * IQR of the hinge.

Overall, > 36% of all proteins involved in ribosome biogenesis, as described in KEGG (29), showed significantly increased expression within the Low XIST RNA population (Fig. 4b). Remarkably >70% of genes encoding proteins involved in cytoplasmic ribosome biogenesis showed increased protein expression (Fig. 4d & e). Specifically, the atypical RIO kinases RIOK1 and RIOK2, which are essential for 40S maturation (30) and SBDS, LSG1 and SPATA5 (a human homologue of yeast Drg1), which are all involved in the final step of 60S maturation, showed significantly increased protein expression within the Low XIST RNA population (Fig. 4e & f). Interestingly, these proteins are involved in cytoplasmic quality control of ribosomes (31–33).

### Proteome specific changes affecting ribosomes and translation initiation

Cytoplasmic ribosomal proteins showed a mean increase of ~13% (p-value 0.001941) in copy number in the Low XIST population (Fig. 5a). Furthermore, the changes in ribosomal protein levels were not uniform between proteins that are part of the 60S and the 40S subunits. The largest increase in the Low XIST population affected 60S ribosomal subunit proteins (Fig. 5b), resulting in a significant change (p-value = 0.0053) in the ratio of protein copy numbers between 60S and 40S ribosomal proteins (Fig. 5c). Overall, ~42% of ribosomal proteins and ribosomal protein S6 kinases were significantly increased in expression in the Low XIST RNA population (Fig. 5d). Interestingly, the most significantly changed protein (p-value <6.87e-07) was p90 ribosomal S6 kinase (RPS6KA3; also significantly increased in the RNAseq data), which is encoded on the X chromosome and is linked to cell growth via increased cap dependent translation through phosphorylation of RPS6 (34) and RPTOR (35) (for all significantly increased X-linked kinases see supplemental figure 2).

**Figure 5.**
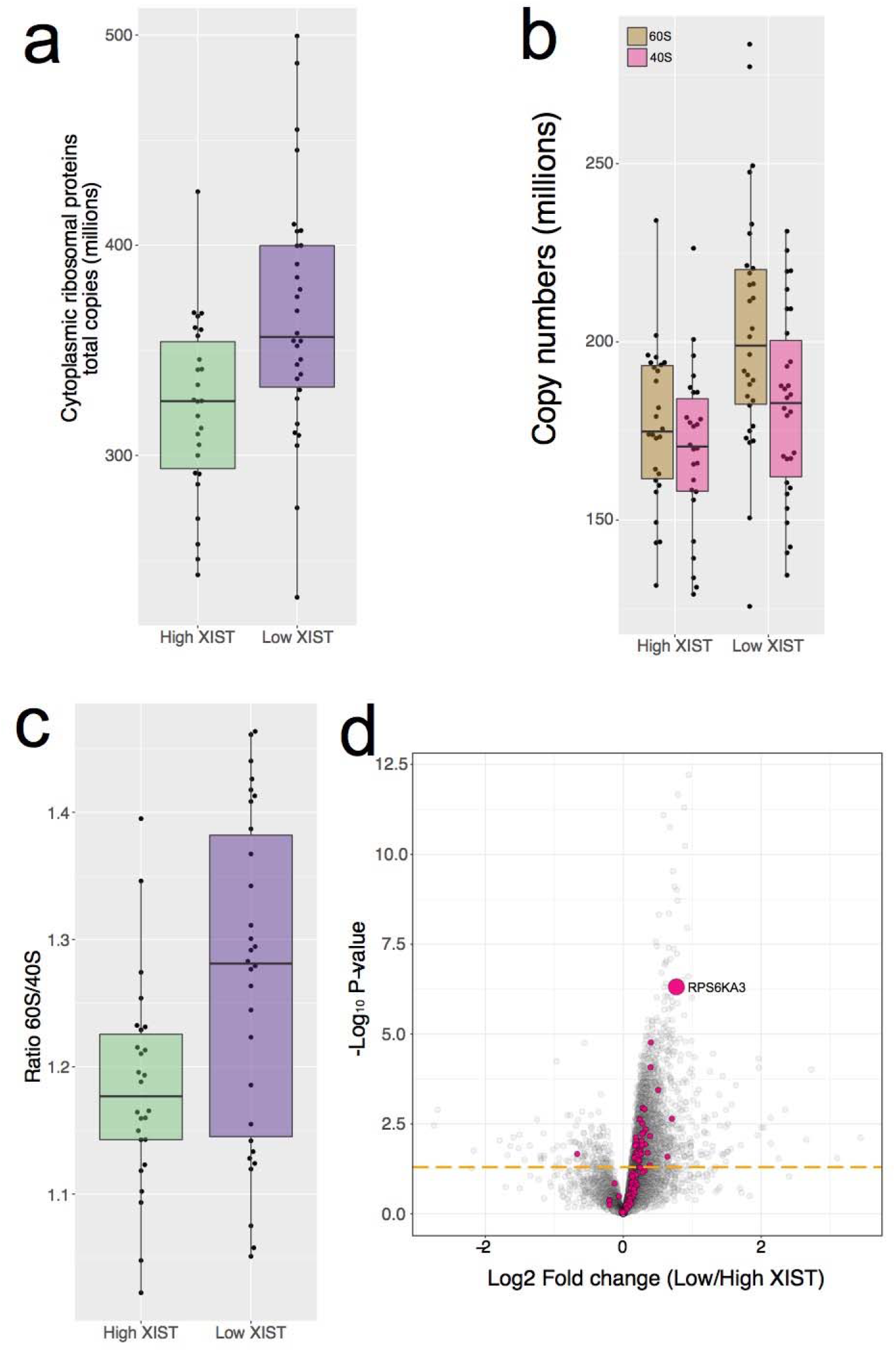
Ribosomes and translational initiation: **(a)** Box plot for the High and Low XIST populations showing the copy numbers for the sum of all cytoplasmic ribosomal proteins. The lower and upper hinges represent the 1st and 3rd quartiles. The upper whisker extends from the hinge to the largest value no further than 1.5 * IQR from the hinge, the lower whisker extends from the hinge to the smallest value at most 1.5 * IQR of the hinge. **(b)** Box plot for the High and Low XIST populations showing the copy numbers for the sum of all 60S (large ribosomal subunit) and 40S (small ribosomal subunit) proteins. The lower and upper hinges represent the 1st and 3rd quartiles. The upper whisker extends from the hinge to the largest value no further than 1.5 * IQR from the hinge, the lower whisker extends from the hinge to the smallest value at most 1.5 * IQR of the hinge. **(c)** Box plot for the High and Low XIST populations showing the ratio of the sum of all 60S to 40S proteins. The lower and upper hinges represent the 1st and 3rd quartiles. The upper whisker extends from the hinge to the largest value no further than 1.5 * IQR from the hinge, the lower whisker extends from the hinge to the smallest value at most 1.5 * IQR of the hinge. **(d)** Volcano plot showing the protein log_2_ fold change between the Low and High XIST populations on the X axis, with the −log_10_ p-value on the y axis. Ribosomal proteins and ribosomal S6 kinases are highlighted in pink; all other proteins are coloured grey. All proteins above the orange line have a p-value lower than 0.05

We also detected a significant increase in expression of the translation initiation factors EIF2S3 and EIF1AX in the Low XIST RNA population. Both EIF1AX and EIF2S3 are part of the 43S preinitiation complex (36), are encoded on the X chromosome and are significantly increased in expression at both the protein and transcript levels (Fig. 6a & b). EIF2S3 is a member of the heterotrimeric eIF2 complex, which delivers an initiator methionyl transfer RNA to the ribosome. Based on protein expression data EIF2S3 appears to be the rate limiting subunit of this complex within the iPSC lines (supplemental figure 3). Similarly, EIF1AX is involved in virtually all the steps from the pre-initiation to ribosomal subunit joining (37), and shows a dramatic median increase of over 2.3 million copies in the low XIST RNA population.

**Figure 6.**
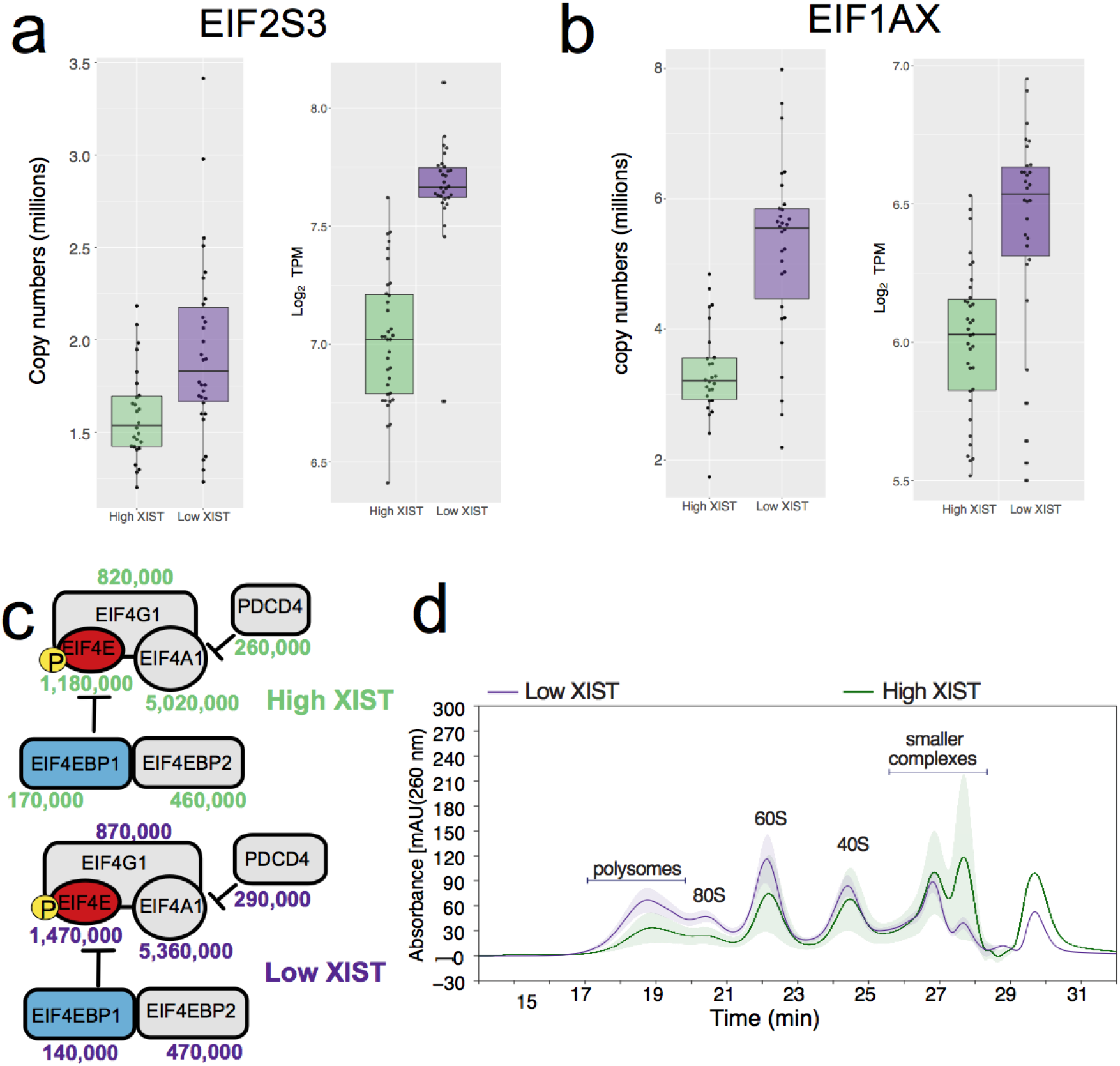
Ribosomes: **(a)** RNA and protein level box plots for the High and Low XIST populations showing the protein copy numbers and Log_2_ TPM for EIF2S3. The lower and upper hinges represent the 1st and 3rd quartiles. The upper whisker extends from the hinge to the largest value no further than 1.5 * IQR from the hinge, the lower whisker extends from the hinge to the smallest value at most 1.5 * IQR of the hinge. **(b)** RNA and protein level box plots for the High and Low XIST populations showing the protein copy numbers and Log_2_ TPM for EIF1AX. The lower and upper hinges represent the 1st and 3rd quartiles. The upper whisker extends from the hinge to the largest value no further than 1.5 * IQR from the hinge, the lower whisker extends from the hinge to the smallest value at most 1.5 * IQR of the hinge. **(c)** Schematic showing the protein copy numbers for the eIF4F complex and it’s inhibitors displayed for both the Low and High XIST populations. Proteins represented via red boxes are significantly increased, light blue boxes are significantly decreased and elements in grey boxes remain unchanged. **(d)** Ribo Mega-sec derived line plot showing the mean Low and High XIST profile with the coloured ribbon representing the standard deviation.

Furthermore, the potential impact of eroded XCI on translation initiation is not limited to genes on the X chromosome. eIF4F is another heterotrimeric protein complex, composed of protein subunits alpha (EIF4A1 & EIF4A3), epsilon (EIF4E) and gamma (EIF4G1 & EIF4G3) (38), which is vital for translation initiation. The stoichiometry of EIF4E, the cap binding subunit, to its inhibitors EIF4EBP1 and EIF4EBP2 (39), is potentially rate limiting within the iPSC lines analysed. Interestingly, in the Low XIST RNA population there is a significant increase in EIF4E levels alongside a parallel significant decrease in EIF4EBP1 (Fig. 6c). The decrease in EIF4EBP1 is particularly relevant in this case as only ~1.2% of all proteins are significantly decreased in expression in the Low XIST RNA population.

In summary, the data are consistent with erosion of XCI in the Low XIST RNA population causing a major change in the proteome that results in a global increase in total protein levels and suggest that this is mediated, at least in part, via increased levels of translation. To test the hypothesis that the iPSC lines with low levels of XIST RNA have increased levels of translation, we compared polysome and ribosome levels between iPSC lines showing either High, or Low levels of XIST RNA, using the Ribo Mega-SEC method (40). This size exclusion chromatography (SEC) based method can separate large protein complexes, including polysomes and ribosomes.

The Ribo Mega-SEC data were remarkably consistent with the proteomics data. This showed that both the polysome containing fractions and 80S containing fractions are increased significantly in extracts from iPSC lines with low levels of XIST RNA, compared to extracts from iPSC lines with high XIST RNA levels (Fig. 6d). Furthermore, the SEC data show a more pronounced increase within the Low XIST RNA lines for 60S, as compared with 40S containing fractions, consistent with the differences seen in ribosomal protein levels in the MS data (Fig. 5b & c). The polysome fractionation data are thus consistent with a model in which erosion of XCI in the iPSC lines expressing low levels of XIST RNA upregulates protein translation capacity and leads to a global increase in protein levels from multiple genes on both autosomes and the X chromosome.

## Discussion

This study provides the first in depth global analysis by mass spectrometry-based proteomics of how erosion of X chromosome inactivation (XCI) affects gene expression at the protein level in human cells. We analysed RNA expression in 74 independent human iPSC lines derived from healthy female donors that were generated by the HipSci consortium (11). This showed that ~40% of these lines expressed very low levels of the long non-coding XIST RNA. Further analysis of bi-allelic expression from X chromosome encoded genes showed that low XIST RNA levels strongly correlated with erosion of XCI. We therefore characterised how the erosion of XCI remodels the human proteome by comparing in detail the levels of RNA and protein expression across 56 female iPSC lines stratified according to high vs low XIST RNA levels.

First, the data show that female derived iPSC lines with low XIST RNA expression have significantly upregulated levels of both RNA and protein from genes on the X chromosome. Including the evidence from increased biallelic expression in the low XIST lines, these data indicate erosion from XCI causes increased protein expression from X linked genes primarily via a transcriptional mechanism.

Second, we also detected a significant increase in protein expression levels from ~26% of genes on autosomes in the low XIST population. However, in contrast with genes on the X chromosome, for most autosomal genes increased protein expression did not correlate with increased RNA expression. Thus, autosome encoded genes showed a low overall RNA:protein fold change correlation (median Pearson correlation for autosomes of 0.27, compared with 0.56 for X). These data indicate that XCI erosion in human cells can affect protein levels encoded by a much wider range of genes than was previously shown by RNAseq data alone, including autosome-linked genes and disease loci.

Third, our comparison of high vs low XIST populations showed that the low XIST expressing cell lines had a median increase of ~13% in total protein content. In considering the potential mechanism causing this increased protein content, several lines of evidence suggest it may result from post transcriptional regulation affecting translation efficiency. Focussing on the proteins that show significant RNA-independent increases in expression in Low XIST lines revealed enrichment for GO terms associated with ribonucleoprotein complex biogenesis. Furthermore, polysome profiling analyses in iPSC lines using the Ribo Mega-SEC method(40) also showed a significant increase in polysomes and 80S ribosomes that correlated with low XIST RNA levels. Thus, both the independent MS proteomics and polysome analyses support the view that cells showing XCI erosion show increased protein translation activity, resulting in a global increase in total protein levels.

In light of the elevated levels of polysomes and increased protein expression observed in the Low XIST iPSC population, it is interesting that the two X-linked genes that encode important regulators of translational initiation, i.e. eIF1AX and eIF2S3, show highly increased expression at both the transcript and protein level. It has been proposed that protein synthesis is principally regulated at the initiation stage(36). The eIF1AX and eIF2S3 proteins are thus candidates for mediating, at least in part, the mechanism whereby erosion of XCI causes an increase in protein translation capacity. Furthermore, both the EIF1AX and EIF2S3 genes have been categorized previously as facultative XCI escapees(41, 42), meaning they are amongst a subset of X-linked genes that can escape repression, despite the global XCI state. This local increased gene dosage effect suggests that even female lines with normal XCI may differ in translational capacity from male cells with only a single X chromosome. Future studies are planned to characterise in detail the differences in gene expression and translation activity between iPSC lines derived from healthy female and male donors.

Our data show that erosion of XCI in human cells has the potential to cause major changes at the level of protein expression, which in turn could have important implications for disease progression and response to therapy in females. The potential clinical relevance is amplified by our finding that expression of many autosomal genes also respond to erosion of XCI at the protein level, with many of the significantly increased proteins encoded on autosomes, such as ERK2(43), FYN(44) and CDK6(45), linked to cancer and other diseases. It will be interesting to analyse this further in future studies to determine how commonly clones may arise in female patients showing significant levels of XCI. It will also be of interest to analyse whether, or to what extent, loss of XIST RNA expression and erosion of XCI may occur during iPSC reprogramming and to determine whether cells with low XIST RNA levels could show a growth advantage during reprogramming due to the changes caused in protein levels.

## Methods

### Generation of iPSC lines

All lines included in this study are part of the HipSci resource and were reprogrammed from primary fibroblasts as previously described(11). Out of the total of more than 800 iPSC lines available within the HipSci resource (www.hipsci.org), 217 lines, predominantly from healthy donors, were selected for in depth proteomic analysis in this study using Tandem Mass Tag Mass Spectrometry. A subset of 74 iPSC lines derived from healthy female donors were then used for the XCI analysis.

### TMT Sample preparation

For protein extraction, iPSC cell pellets were washed with ice cold PBS and redissolved immediately in 200 μL of lysis buffer (8 M urea in 100 mM triethyl ammonium bicarbonate (TEAB)) and mixed at room temperature for 15 minutes. The DNA content of the cells was sheared using ultrasonication (6 X 20 s on ice). The proteins were reduced using tris-carboxyethylphosphine TCEP (25 mM) for 30 minutes at room temperature, then alkylated in the dark for 30 minutes using iodoacetamide (50 mM). Total protein was quantified using the EZQ assay (Life Technologies). The lysates were diluted with 100 mM TEAB 4-fold for the first digestion with mass spectrometry grade lysyl endopeptidase, Lys-C (Wako, Japan), then further diluted 2.5-fold before a second digestion with trypsin. Lys-C and trypsin were used at an enzyme to substrate ratio of 1:50 (w/w). The digestions were carried out overnight at 37°C, then stopped by acidification with trifluoroacetic acid (TFA) to a final concentration of 1% (v:v). Peptides were desalted using C18 Sep-Pak cartridges (Waters) following manufacturer’s instructions.

For tandem mass tag (TMT)-based quantification, the dried peptides were re-dissolved in 100 mM TEAB (50 ml) and their concentration was measured using a fluorescent assay (CBQCA, Life Technologies). 100 mg of peptides from each cell line to be compared, in 100 ml of TEAB, were labelled with a different TMT tag (20 mg mb^1^ in 40 ml acetonitrile) (Thermo Scientific), for 2 h at room temperature. After incubation, the labelling reaction was quenched using 8 ml of 5% hydroxylamine (Pierce) for 30 min and the different cell lines/tags were mixed and dried *in vacuo*.

The TMT samples were fractionated using off-line high-pH reverse-phase (RP) chromatography: samples were loaded onto a 4.6 × 250 mm Xbridge BEH130 C18 column with 3.5-mm particles (Waters). Using a Dionex bioRS system, the samples were separated using a 25-min multistep gradient of solvents A (10 mM formate at pH 9) and B (10 mM ammonium formate pH 9 in 80% acetonitrile), at a flow rate of 1 ml min^-1^. Peptides were separated into 48 fractions, which were consolidated into 24 fractions. The fractions were subsequently dried and the peptides re-dissolved in 5% formic acid and analysed by LC–MS/MS.

### TMT LC–MS/MS

*TMT-based analysis*. Samples were analysed using an Orbitrap Fusion Tribrid mass spectrometer (Thermo Scientific), equipped with a Dionex ultra-high-pressure liquid-chromatography system (RSLCnano). RPLC was performed using a Dionex RSLCnano HPLC (Thermo Scientific). Peptides were injected onto a 75 μm × 2 cm PepMap-C18 pre-column and resolved on a 75 μm × 50 cm RP-C18 EASY-Spray temperature-controlled integrated column-emitter (Thermo Scientific), using a four-hour multistep gradient from 5% B to 35% B with a constant flow of 200 nl min^-1^. The mobile phases were: 2% ACN incorporating 0.1% FA (solvent A) and 80% ACN incorporating 0.1% FA (solvent B). The spray was initiated by applying 2.5 kV to the EASY-Spray emitter and the data were acquired under the control of Xcalibur software in a data-dependent mode using top speed and 4 s duration per cycle. The survey scan was acquired in the orbitrap covering the *m/z* range from 400 to 1,400 Thomson with a mass resolution of 120,000 and an automatic gain control (AGC) target of 2.0 × 10^5^ ions. The most intense ions were selected for fragmentation using CID in the ion trap with 30% CID collision energy and an isolation window of 1.6 Th. The AGC target was set to 1.0 × 10^4^ with a maximum injection time of 70 ms and a dynamic exclusion of 80 s.

During the MS3 analysis for more accurate TMT quantifications, 5 fragment ions were co-isolated using synchronous precursor selection using a window of 2 Th and further fragmented using HCD collision energy of 55%. The fragments were then analysed in the orbitrap with a resolution of 60,000. The AGC target was set to 1.0 × 10^5^ and the maximum injection time was set to 105 ms.

### Ribo Mega-SEC iPSC lines

For the Ribo Mega-SEC analyses 4 iPSC lines with High XIST RNA levels (‘iiyk_2’, ‘iiyk_4’, ‘nufh_3’ and nufh_4’) and 3 lines with Low XIST RNA levels (‘fawm_4’, ‘bawa_1’ and ‘aizi_3’) were used.

### Ribo Mega-SEC

Ribo Mega-SEC for the separation of polysomes and ribosomal subunits using size exclusion chromatography was performed as previously reported (40), with a slight modification. Briefly, 2.5 × 10^6^ cells were washed once with ice-cold PBS, scraped in ice-cold PBS and collected by centrifugation at 500 g for 5 min (all centrifugations at 4°C). The cells were lysed by vortexing for 10 sec in 250 μl of polysome extraction buffer (20 mM Hepes-NaOH (pH 7.4), 130 mM NaCl, 10 mM MgCl2, 5% glycerol, 1% CHAPS, 0.2 mg/ml heparin, 2.5 mM DTT, 20 U SUPERase In RNase inhibitor, cOmplete EDTA-free Protease inhibitor), incubated for 15 min on ice, and centrifuged at 17,000 g for 10 min. Supernatants were filtered through 0.45 μm Ultrafree-MC HV centrifugal filter units (Millipore).

Using a Dionex Ultimate 3,000 Bio-RS uHPLC system (Thermo Fisher Scientific), a SEC column (Agilent Bio SEC-5, 2,000 Å pore size, 7.8 × 300 mm with 5 μm particles) was equilibrated with three column volumes of filtered SEC buffer (20 mM Hepes-NaOH (pH 7.4), 60 mM NaCl, 10 mM MgCl_2_, 0.3% CHAPS, 0.2 mg/ml heparin, 2.5 mM DTT, 5% glycerol) (all column conditioning and separation at 5°C) and 100 μl of 10 mg/ml of filtered bovine serum albumin (BSA) solution diluted by PBS was injected once to block the sites for non-specific interactions. After monitoring the column condition by injecting standards, including 10 μl of 10 mg/ml BSA solution and 5 μl of HyperLadder 1 kb (BIOLINE), 200 μl of the filtered cell lysates was injected onto the pre-equilibrated SEC column. The flow rate was 0.4 ml/min and the chromatogram was monitored by measuring UV absorbance at 215, 260 and 280 nm with a 1 Hz data collection rate by the Diode Array Detector.

### RNA-seq data processing

Raw RNA-seq data were obtained from the ENA project: ERP007111. CRAM files were merged on a sample level and converted to FASTQ format. The reads were trimmed to remove adapters and low quality bases (Trim Galore!), followed by read alignment using STAR(46) (version: 020201), using the two-pass alignment mode and the default parameters as proposed by ENCODE (c.f. STAR manual). All alignments were relative to the GRCh37 reference genome, using ENSEMBL 75 as transcript annotation.

Samples with low quality RNA-seq were discarded if they had either less than 2 billion bases aligned, had less than 30% coding bases, or had a duplication rate higher than 75%. Gene-level RNA expression was quantified from the STAR alignments using featureCounts(47) (v1.6.0), which was applied to the primary alignments using the “-B” and “-C” options in stranded mode, using the ENSEMBL 75 GTF file. Quantifications per sample were merged into an expression table using the following normalization steps. First, gene counts were normalized by gene length. Second, the counts for each sample were normalized by sequencing depth using the edgeR(48) adjustment. The TPM values as returned by Salmon were combined into an expression table.

### Identification & Quantification

The TMT-labelled samples were collected and analysed using Maxquant (49, 50) v. 1.6.3.3. The FDR threshold was set to 1% on the Peptide Spectrum Match (PSM) Protein level. Proteins and peptides were identified using the UniProt human reference proteome database (SwissProt). Run parameters have been deposited to PRIDE(12) along with the full MaxQuant quantification output (PDX010557).

Data for the analysis were obtained from the ProteinGroups.txt output of MaxQuant. Contaminants, reverse hits and ‘only identified by site’ proteins were excluded from analysis. Overall, we quantified 9,631 protein groups in at least one of the samples.

### Protein filtering

For additional stringency and to reduce batch variation, only proteins with 3 or more ‘Unique + razor peptides’ were considered.

### iPSC Copy number generation

Protein copy numbers were calculated using the proteomic ruler (14) and using the MS3 reporter intensity. An additional batch correction factor for each TMT experiment was applied as described here (15).

### Protein content

The protein content for both the High and Low XIST RNA populations was calculated based on the copy numbers for each line and then converted into picograms. The molecular weight of each protein was multiplied by the number of copies for the corresponding protein and this was then summed for all proteins to calculate the protein content.

### Chromosome mapping

To map gene products to their specific chromosomes, we utilised the UniProt (51) protein-chromosome mapping service. We used their output to produce a list of unique proteins for each specific chromosome. Subsequently, we mapped the proteins detected in our iPSC dataset to their corresponding chromosomes based on the UniProt mapping file.

### X-inactivation stratification and analysis

Based on the RNAseq data, 74 iPSC lines were classified into 3 distinct categories based on XIST expression. 30 iPSC lines where XIST expression was <1 Log_2_ TPM were classified as ‘Low XIST’. 35 where XIST expression was higher than 2.75 Log_2_ TPM were classified as High XIST and the remaining 9 lines were classified as ‘Medium’ XIST.

### High XIST filtering

The High XIST population contained two proteomic experiments, PT7422 and PT6386, contributing a large number of High XIST replicates within their 10-plex TMT experiment. As the maximum number of replicates per 10-plex within the Low XIST group was 4, we performed hierarchical clustering to reduce the number of a lines contributed by PT7422 and PT6386 to a maximum of 4 in order to minimise batch effects. The final number of lines with High XIST post filtering was 26.

### Enrichment analysis

All of the enrichment analyses were done using Panther(28) and with the 8,511 proteins which were detected in both the RNAseq and proteomics datasets as background. We performed a biological process overrepresentation analysis for all proteins that were significantly increased (q-value <0.05), where the corresponding transcript was not significantly increased in expression. Furthermore, an additional biological process overrepresentation of significantly increased (q-value<0.05) ribosome biogenesis proteins was carried out.

### 60S/40S ratio

The ratios were calculated by summing the copy numbers from all proteins of the 60S ribosomal subunit divided by the sum of all copy numbers from the 40S ribosomal subunit, for each individual iPSC line.

### Differential expression analysis

Fold changes and P-values were calculated in R utilising the bioconductor package LIMMA(52) version 3.7. The Q-values provided were generated in R using the “qvalue” package version 2.10.0.

### UniProt to Ensembl mapping

Mapping of UniProt accessions to Ensembl gene identifiers was done in R using the “UniProt.ws” package version 2.24.1

### Kinase map

The kinase map was generated within the Encyclopedia of Proteome Dynamics (13) using the KinoViewer (53).

## Acknowledgements

We thank all of our collaborators who worked on the HipSci project. We would also like to thank D. Cantrell, M. Stavridis, G. Findlay, J. Marchingo and F. Dossin for all the insightful discussions as well as L. Davidson within the University of Dundee Stem Cell Facility. We would also like to thank all members of the Lamond Laboratory.

## Data and materials availability

All of the mass-spectrometry raw files and the MaxQuant outputs have been uploaded to PRIDE and are publicly available under accession PXD10557. The RNAseq data has been uploaded to the ENA project are publicly available under accession PRJEB7388.

## Author contributions

A.J.B. conceived the study. A.I.L. & O.S supervised the project. H.Y assisted in the interpretation of the data as well as designing and executing the polysome profiling experiments. H.J & J.H. assisted in the interpretation of the data. D.B. designed and performed all proteomics experiments. D.S. and B.M. processed and prepared all RNAseq data. D.S performed the allele specific analysis. A.J.B. performed the data analysis and interpretation. The paper was written by A.J.B. and A.I.L and edited by all authors.

